# CROP: A CRISPR/Cas9 guide selection program based on mapping guide variants

**DOI:** 10.1101/2020.03.05.979880

**Authors:** Victor Aprilyanto, Redi Aditama, Zulfikar Achmad Tanjung, Condro Utomo, Tony Liwang

## Abstract

The off-target effect, in which DNA cleavage was conducted outside the targeted region, is a major problem which limits the applications of CRISPR/Cas9 genome editing system. CRISPR Off-target Predictor (CROP) is standalone program developed to address this problem by predicting off-target propensity of guide RNAs and thereby allowing the user to select the optimum guides. The approach used by CROP involves generating substitution, deletion and insertion combinations which are then mapped into the reference genome. Based on these mapped variants, scoring and alignment are conducted and then reported as a table comprising the off-target propensity of all guide RNAs from a given gene sequence. The Python script for this program is freely available from: https://github.com/vaprilyanto/crop.

## Introduction

Clustered regularly interspaced short palindromic repeats (CRISPR) is a versatile genome editing tool which relies on the activity of CRISPR-associated (Cas) enzymes to break the DNA strand guided by short 20-nucleotide RNA sequence also known as a guide^1–4^. This RNA-guided nuclease (RGN) system has been widely used to knockdown or edit genomic locations across various organisms, notably human, animals, and crop plants. However, this system has one caveat in which it gives off-target cleavage, resulting double-strand break (DSB) at locations other than targeted ones^5,6^.

A number of efforts has been conducted to minimalize this off-target effects, ranging from designing high fidelity Cas9^7–9^ to selecting high specificity guide RNAs (gRNA)^2,10,11^. In the latter approach, the specificity of 20-nucleotide gRNA could be predicted based from the base types and positions in the guide sequence^2^. This observation leads to a number of off-target prediction webserver programs, such as CCTop^12^, Cas-OFFinder^13^, CRISPOR^14^, CRISPR-PLANT^15^, CRISPR-P^16^ and others which are widely used to select gRNA with high specificity. However, the availability of the reference genomes in those webservers often restricts one to predict the off-target of a given guide at organism which genome is unavailable. In this study, we introduced CRISPR Off-target Predictor (CROP), a program for guide RNA off-target prediction which allows the user to use own’s genome of interest. The approach used in CROP is by creating guide variants via substitution, deletion or insertion schemes up to four positions along the twenty nucleotide-long gRNA sequence. This approach is designed to simulate DNA:RNA heteroduplex pairing with mismatch, RNA and DNA bulges. The resulting variants from each gRNA will subsequently be mapped to the reference genome, in which the mapped variants are then used to estimate the off-target propensity score of the corresponding gRNA.

## Materials and Methods

### Generating Guide Variants and Mapping to Genome

All the steps conducted in CROP are summarized in Figure 1. For guide finding step, CROP lists all the available 20-nucleotide guide sequences from the input DNA sequence. Then in variant generation step, a number of variants were generated from each guide. This step was carried out through generating all possible positional combinations ranging from one up to four nucleotides across guide’s length. These positional combinations were then used in three subsequent schemes of guide modifications, namely substitution, deletion and insertion (Figure 1). In substitution scheme guide variants were created by replacing bases at positions given by the positional combinations into all possible combinations of DNA bases. In deletion scheme, the bases at these positions were deleted, giving shortened variants from the original guide. The last scheme uses these positional combinations to insert bases into the guide sequence to yield longer variants as the result. In addition to base modifications inside the 20-nucleotide guide sequence, eight combinations of NRG protospacer adjacent motif (PAM) consisting of AGG, TGG, GGG, CGG, AAG, TAG, GAG, and CAG were added as prerequisites for *Streptococcus pyogenes* Cas9 (SpCas9) cleavage, therefore multiplies the total variants to eight fold. All these variants were then mapped towards the reference genome using Bowtie read-mapping^17^ using zero mismatches (`-v 0`) and reporting all available mapped locations per variant (`-a`). The mapping-result was reported as a sequence alignment map (.sam)-formatted file which serves as the base for further scoring and alignment processes.

**Figure 1.**
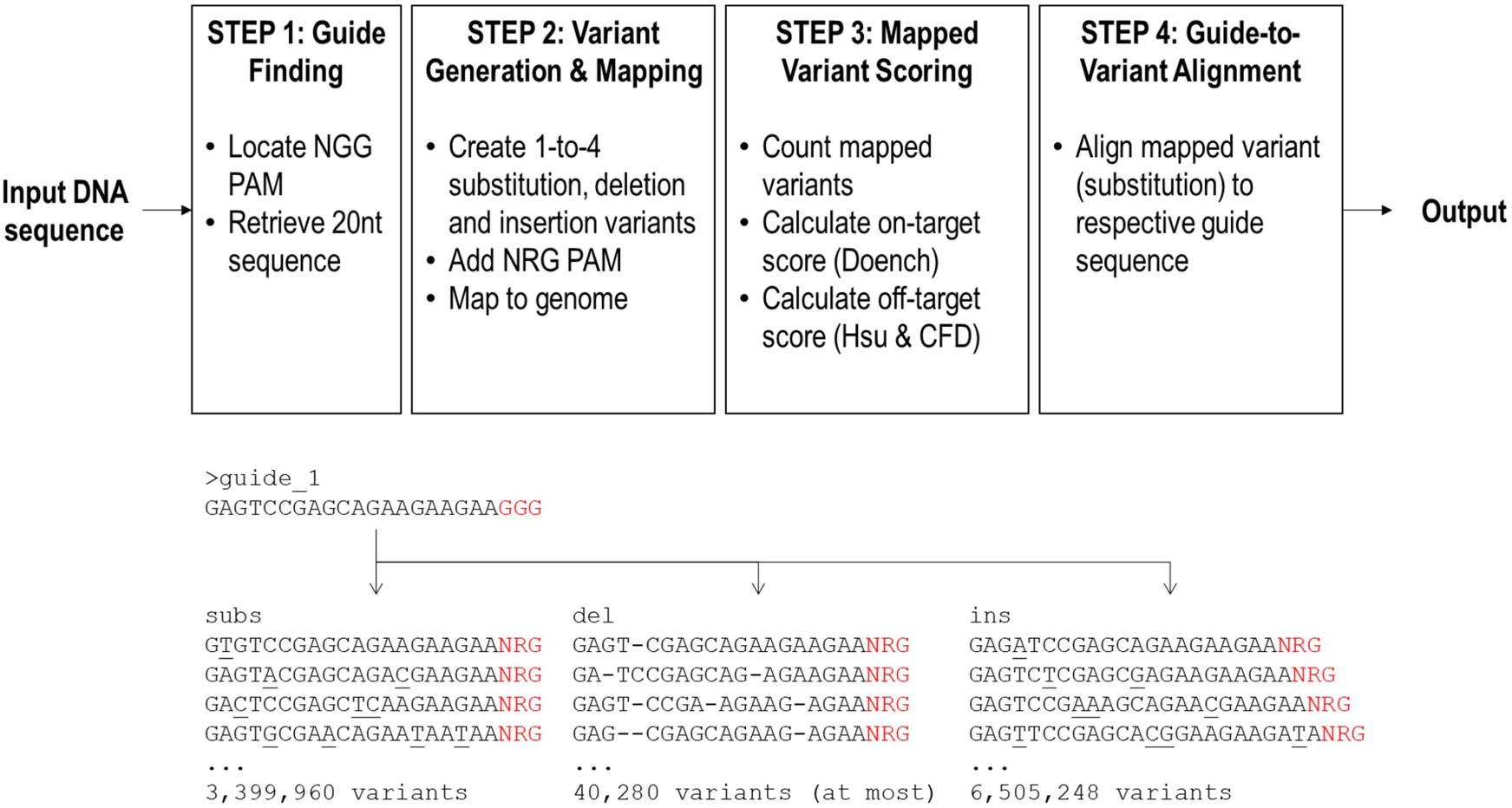
CROP working scheme (top) and generation of variants from a 20-nucleotide guide sequence (bottom).

### Scoring Functions, Alignment and Parallelization

To assess off-target propensity CROP uses counts mapped variants along with Hsu^2^ and CFD^11^ off-target scoring calculations. The mapped variant counts included mapped substitution, deletion and insertion variants. The python scripts for both scoring systems were adopted from CRISPOR program^18^ through GitHub (https://github.com/maximilianh/crisporWebsite) with minor modifications including code simplifications and CFD score re-scaling. Using only mapped substitution count, Hsu and CFD off-target scores were then calculated for each guide-variant pair. The usage of only mapped substitution was due to the requirement of the same sequence length between guide and variants. After that, cumulative Hsu and CFD scores among all guide-variant pairs for one guide was calculated according to each aggregate scores^16,19^.

CROP also included on-target scores for each guide, including Doench score^10^ as well as 30-nucleotide sequence bearing N_4_-N_20_-NGG-N_3_ (N_20_ = 20nt guide sequence) format for calculating Azimuth score through Machine learning-based end-to-end CRISPR/Cas9 guide design (https://crispr.ml/) website^11,20^. In addition to scoring, the program also aligned all the mapped variants to the original guide sequence and marked mismatches, highlighting the mismatches for all pairs. In order to automate all the process, a linux script (core.sh) was written to develop a pipeline which consists of four steps, namely: (i) finding guide sequences, (ii) creating guide variants and mapping to reference genome, (iii) variants scoring and (iv) variant alignment. Another Linux script (crop.sh) was also written as the main command file to allow running multiple core.sh in parallel, thereby allowing the user to conduct multiple CRISPR guide analyses in a single operation.

### Test Cases

In order to test its performance, CROP was tested for predicting optimum guides for targeting palmitoyl-acyl carrier protein thioesterase gene (PATE; GenBank accession number XM_010926998) in African oil palm (*Elaeis guineensis*). The oil palm genome (assembly EG5) was downloaded from NCBI Genome database and used as the reference genome for mapping.

## Results and Discussion

### Total Combination of a Guide’s Unique Variants

In order to list all the possible off-targets bearing few substitutions, deletions or insertions for one guide, CROP produces all possible positional combinations of the guide. Under substitution scheme the positional combination took place across the whole guide’s length, where the bases at these positions were substituted into another bases, creating a total of unique substitution variants *S*_U_ according to:

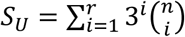

where 1 ≤ *r* ≤ *n* is the number of substituted position and *n* is the number of nucleotides subjected for substitution. For a 20-nucleotide long, this substitution scheme will create 424,996 unique variants. Since this scheme generates variants with sequence length equal to the original guide, the sets of variants are nested, with set generated through lower *r* value is a subset of higher *r* value. This observation was then used to modify the corresponding script by only generating variants from the highest *r* value, which was 4.

In the deletion scheme the deletion was carried out towards 19 out of 20 nucleotides along the guide’s sequence, avoiding 5’-end nucleotide deletion which will create 19-nucleotide subsequence identical to the original guide. The total unique variants generated from this deletion scheme (*D*_*U*_) can be calculated according to:

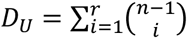

where 1 ≤ *r* ≤ *n* is the number of substituted position and (*n* − 1) is the number of nucleotides subjected for deletion. However, the above formula only provides the maximum number of unique deletion variants which could be generated without specifying the exact number. Using the above formula, at most 5,035 unique variants could be generated from a 20-nucleotide guide sequence. The exact number of unique variants was difficult to calculate since different sequences yields different unique deletion variants. It is thought that the occurrence of *r*-nucleotide repeats affect this number. As for example, *D*_*U*_ for a guide comprising solely of A’s is 4, while guide AAAAATTTTTGGGGGCCCCC and ATGCGTCATCAGACGTAGTC would give *D*_*U*_ 69 and 4647, respectively.

Then in the insertion scheme, a combination of four nucleotides were inserted in positions between the two nucleotides in guide sequence plus one position between the guide’s 3’-end and the first PAM site, giving a total of 20 sites available for insertion. The total unique variants generated from this scheme

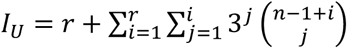

where 1 ≤ *r* ≤ *n* is the number of substituted position and (*n* − 1 + *i*) is the number of sites subjected for insertion. In opposite to the deletion scheme, the variants generated in this insertion scheme had longer sequences. The total unique combination under this scheme was also larger than substitution and deletion schemes as was calculated by the formula. The total unique insertion variants generated from one 20-nucleotide guide was 813,160, comprising of 61; 1,954; 43,726 and 767,419 for 1, 2, 3 and 4 insertions, respectively.

In total, these three schemes would give 1,243,190 unique variants from one guide. In addition to NGG PAM, CROP also considers NAG PAM which is reported to yield Cas9 cleavage albeit with lower probability^2,10^. Therefore, with a combination of NRG PAM (R = A or G), the total unique variants would increase eight folds to 9,945,520.

### CROP Runtime and Output Delivery

Using a standard desktop computer (i5, 8 Gb RAM) CROP required more or less 100 seconds from guide finding until guide-to-variant alignment steps (Figure 1). A typical gene fragment giving 100 guide sequences would require around three hours. Although the runtime could be improved using more powerful cores, it was still considered less efficient compared to other web-based CRISPR programs, which usually produces results in minutes. The main limiting factor for CROP runtime was likely in the variant generation step which requires the program to generate almost ten million variants per guide. Several approaches could be attempted to improve CROP’s efficiency, including the usage of higher number of computer cores or considering subsets among the total variants which are frequently observed to cause off-targeting in Cas9.

Using more computer cores could increase the number of parallel processes executed at one time. A standard desktop computer (i5) contains four cores which means that it could predict four guides at one time. A standard workstation with 88 cores could predict 22 times more guides at one time compared to the standard desktop. On variant subset consideration, several studies on CRISPR applications^1,2,21^ reported that the actual off-targets are locations in which the sequence contain less or no mismatches at the PAM proximal region (ten nuelcotides adjacent to PAM). This region is therefore an important determinant for Cas9 activity. Assuming that a Cas9-gRNA complex would only cleave locations given a perfect match in the PAM proximal, then the variant set could be restricted to only containing sequences with mismatches at PAM distal region (nucleotides 11-20 adjacent to PAM). With such restrictions, the time needed to create guide variants and overall prediction process will be faster

For each queried DNA sequence, CROP give a result in four directories, namely: (i) `align`, containing alignment between the guide sequence with its mapped variants; (ii) `ot_maps`, containing sam-formatted file listing mapped guide variants; (iii) `single_guides`, containing single-fasta files of single-guide RNA which could be used for mFold RNA loop prediction^22^; and (iv) `outfiles`, containing score table, raw sequence, and multi-fasta files. The guides in raw sequence file are written by including additional bases in N_4_-guide-NGG-N_3_ format as input for Azimuth score calculation in https://crispr.ml/ website. The score file contained a number of information for each guide, including GC content, Doench on-target score, map counts, Hsu & CFD off-target score, and number of on target count. All on- and off-target scores are set to a 0-100 scale. A log file was also written in the result directory, containing all the activities that CROP carried out towards the corresponding DNA sequence.

### Highly Variable On- and Off-target Scores Among PATE Exon-1 Guides

Among nearly 10 million variants per guide, only a tiny fraction of which mapped to the oil palm EG5 reference genome. For PATE exon-1 gene, this fraction ranged from 0.0003% to 0.01% which were dominated by substitution and deletion variants. CROP produced 79 guides from PATE exon-1 gene, comprising 35 forward and 44 reverse guides (Figure 2; Supplementary Table S1). Based on these counts, the mapped insertion variants (*ins*) had the lowest counts among all three. The mapped substitution (*subs*) and deletion (*del*) dominated the counts with majority of the guides had higher counts on the former. These mapped counts varied widely from as low as zero in mapped insertion variants to near 900 in mapped substitution variants. Similar to off-target counts, the Hsu and CFD off-target scores also displayed wide variations among the guides with the former generally being larger than the latter. Such score difference might be related with the employment of different off-target databases by both scoring system. The database used in CFD calculation was inferred from a larger dataset compared to the Hsu’s^2,11^. This might hint that CFD is a better predictor in regards of off-target propensity of guide RNA^23^.

**Figure 2.**
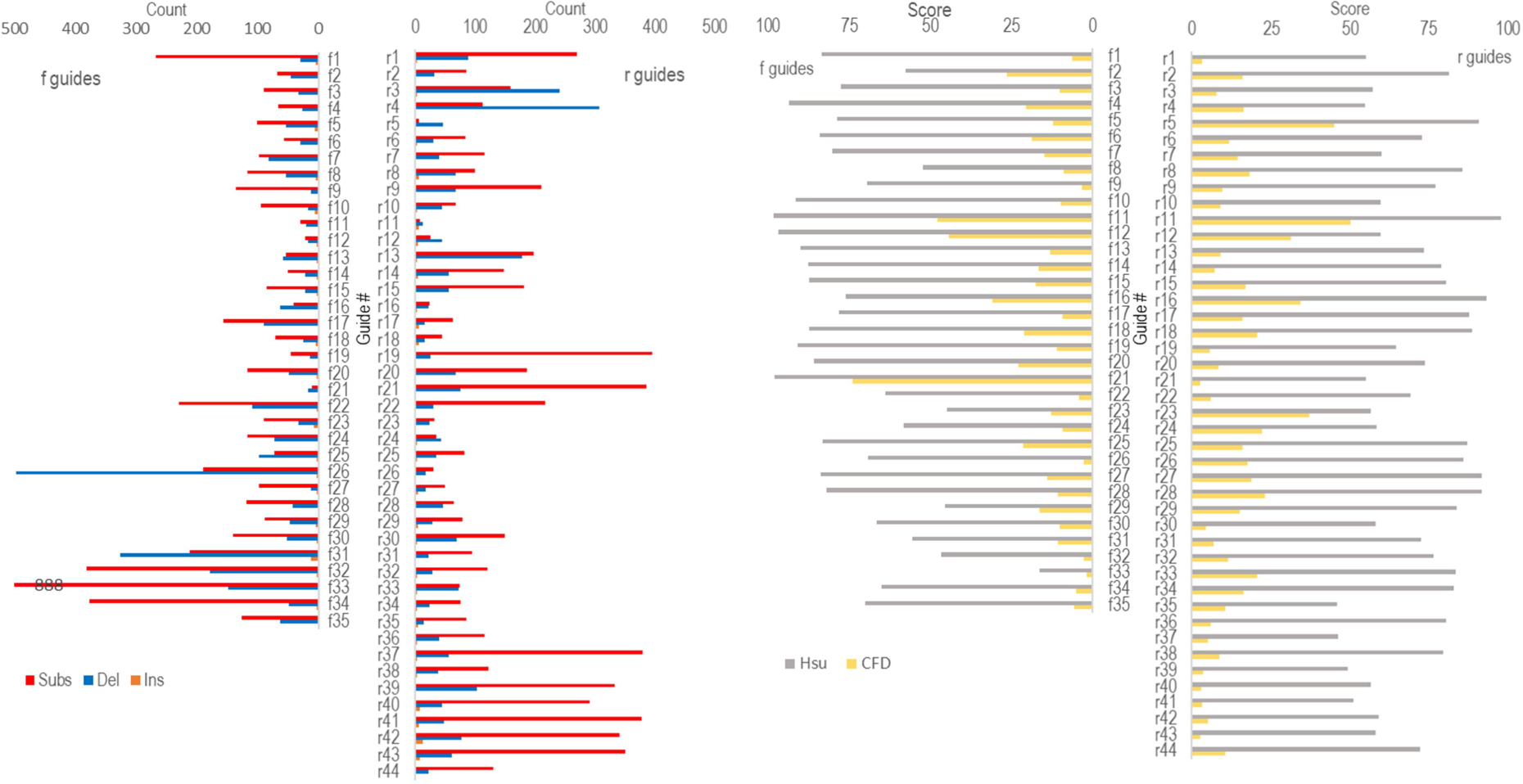
CROP prediction on guides derived from oil palm PATE exon-1 gene (XM_010926998). Left: off-target counts of mapped substitution, deletion and insertion variants; right: off-target Hsu and CFD scores derived from mapped substitution variants. Higher score translates to lower off-target propensity.

Ideally, guide sequences to be used in genome editing applications should be ones which exhibit high on-target while maintaining low off-target capabilities. However, as shown in the case of PATE exon-1, an opposite relationship was observed between on-target (Doench^11^) and off-target (Hsu^2^ and CFD^11^) scores. Guides possessing high off-target scores, those that possess high specificity, tend to give poor on-target scores and vice versa. Such opposite relationship tends to limit the search for optimal guides and therefore a compromise has to be made. For example, guide r5 (Figure 2; Supplementary Table S1) was considered to be the best among all 79 guides since it possess a high on-target and adequate off-target scores. Other guide such as guide f21 also possessed high off-target scores but scored poorly on on-target score. To avoid inability of Cas9 cleavage cause by such guide, an *in vitro* Cas9 test is recommended for all the selected guides prior to *in vivo* applications.

## Conclusion

CROP is a simple and flexible standalone program which predicts off-target propensity of guide RNAs from a gene via mapping possible guide variants towards the genome. A test using oil palm PATE exon-1 gene revealed that majority of the guides possess high off-target propensity, thus they are not recommended to be used in CRISPR genome editing applications.

## Acknowledgements

This work was supported by the management of PT SMART Tbk. We thank Reno Tryono, Roberdi and Yudistira Wahyu Kurnia for proof-reading this manuscript.

## References

1. Jinek, M., Chylinski, K., Fonfara, I., Hauer, M. & Doudna, J. A. A programmable dual-RNA-guided DNA endonuclease in adaptive bacterial immunity. Science (80-.). 337, 816–821 (2012).

2. Hsu, P. D. et al. DNA targeting specificity of RNA-guided Cas9 nucleases. Nat. Biotechnol. 31, 827–832 (2013).

3. Sternberg, S. H., Redding, S., Jinek, M., Greene, E. C. & Doudna, J. A. DNA interrogation by the CRISPR RNA-guided endonuclease Cas9. Nature 507, 62–67 (2014).

4. Barrangou, R. & Doudna, J. A. Applications of CRISPR technologies in research and beyond. Nat. Biotechnol. 34, 933–941 (2016).

5. Wu, X., Kriz, A. J. & Sharp, P. A. Target specificity of the CRISPR-Cas9 system. Quant. Biol. 2, 59–70 (2014).

6. Tsai, S. Q. & Joung, J. K. Defining and improving the genome-wide specificities of CRISPR–Cas9 nucleases. Nat. Rev. Genet. 17, 300–312 (2016).

7. Slaymaker, I. M. et al. Rationally engineered Cas9 nucleases with improved specificity. Science (80-.). 351, 84–88 (2016).

8. Kleinstiver, B. P. et al. High-fidelity CRISPR-Cas9 nucleases with no detectable genome-wide off-target effects. Nature 529, 490–495 (2016).

9. Vakulskas, C. A. et al. A high-fidelity Cas9 mutant delivered as a ribonucleoprotein complex enables efficient gene editing in human hematopoietic stem and progenitor cells. Nat. Med. 24, 1216–1224 (2018).

10. Doench, J. G. et al. Rational design of highly active sgRNAs for CRISPR-Cas9-mediated gene inactivation. Nat. Biotechnol. 32, 1262–1267 (2014).

11. Doench, J. G. et al. Optimized sgRNA design to maximize activity and minimize off-target effects of CRISPR-Cas9. Nat. Biotechnol. 34, 184–191 (2016).

12. Stemmer, M., Thumberger, T., Del Sol Keyer, M., Wittbrodt, J. & Mateo, J. L. CCTop: An intuitive, flexible and reliable CRISPR/Cas9 target prediction tool. PLoS One 10, 1–11 (2015).

13. Bae, S., Park, J. & Kim, J. S. Cas-OFFinder: A fast and versatile algorithm that searches for potential off-target sites of Cas9 RNA-guided endonucleases. Bioinformatics 30, 1473–1475 (2014).

14. Haeussler, M. et al. Evaluation of off-target and on-target scoring algorithms and integration into the guide RNA selection tool CRISPOR. Genome Biol. 17, 1–12 (2016).

15. Minkenberg, B., Zhang, J., Xie, K. & Yang, Y. CRISPR-PLANT v2: An online resource for highly specific guide RNA spacers based on improved off-target analysis. Plant Biotechnol. J. (2018). doi:10.1111/pbi.13025

16. Lei, Y. et al. CRISPR-P: A web tool for synthetic single-guide RNA design of CRISPR-system in plants. Mol. Plant 7, 1494–1496 (2014).

17. Langmead, B., Trapnell, C., Pop, M. & Salzberg, S. L. Ultrafast and memory-efficient alignment of short DNA sequences to the human genome. Genome Biol. 10, (2009).

18. Haeussler, M. et al. Evaluation of off-target and on-target scoring algorithms and integration into the guide RNA selection tool CRISPOR. Genome Biol. 17, 1–12 (2016).

19. Liu, H. et al. CRISPR-P 2.0: An Improved CRISPR-Cas9 Tool for Genome Editing in Plants. Mol. Plant 10, 530–532 (2017).

20. Listgarten, J. et al. Prediction of off-target activities for the end-to-end design of CRISPR guide RNAs. Nat. Biomed. Eng. 2, 38–47 (2018).

21. Tsai, S. Q. et al. GUIDE-seq enables genome-wide profiling of off-target cleavage by CRISPR-Cas nucleases. Nat. Biotechnol. 33, 187–198 (2015).

22. Zuker, M. Mfold web server for nucleic acid folding and hybridization prediction. Nucleic Acids Res. 31, 3406–3415 (2003).

23. Concordet, J. P. & Haeussler, M. CRISPOR: Intuitive guide selection for CRISPR/Cas9 genome editing experiments and screens. Nucleic Acids Res. 46, W242–W245 (2018).

