## Supplementary Table 1 for "CROP: A CRISPR/Cas9 guide selection program based on mapping guide variants"

**Supplementary Table S1. CROP result on oil palm palmitoyl acyl thioesterase exon 1**

| **Name** | **Seq** | **PAM** | **GC** | **Doench** | **Subs** | **Dels** | **Ins** | **Total** | **Hsu** | **CFD** | **On Tgt** |
| --- | --- | --- | --- | --- | --- | --- | --- | --- | --- | --- | --- |
| patec7-e1_f1 | CACACCATCTTTCTCCCCCA | CGG | 55 | 66.38 | 268 | 30 | 4 | 302 | 83.235 | 6.233 | 1 |
| patec7-e1_f2 | GCAAAAGCTTCGAAGACCAT | TGG | 45 | 67.751 | 68 | 46 | 0 | 114 | 57.703 | 26.462 | 1 |
| patec7-e1_f3 | GCTTCGAAGACCATTGGTGA | AGG | 50 | 29.204 | 90 | 32 | 0 | 122 | 77.28 | 10.297 | 1 |
| patec7-e1_f4 | TGGTGAAGGCTCCGAGAATT | TGG | 50 | 1.865 | 66 | 27 | 1 | 94 | 93.248 | 20.613 | 1 |
| patec7-e1_f5 | GCTCCGAGAATTTGGATGTT | CGG | 45 | 7.374 | 101 | 54 | 6 | 161 | 78.504 | 12.194 | 1 |
| patec7-e1_f6 | CTCCGAGAATTTGGATGTTC | GGG | 45 | 0.708 | 56 | 29 | 1 | 86 | 83.841 | 18.672 | 1 |
| patec7-e1_f7 | TCCGAGAATTTGGATGTTCG | GGG | 45 | 12.315 | 97 | 82 | 1 | 180 | 80.067 | 14.838 | 1 |
| patec7-e1_f8 | AGCCAAACCCACCTCTTCTT | CGG | 50 | 10.796 | 117 | 54 | 4 | 175 | 52.278 | 9.042 | 1 |
| patec7-e1_f9 | CAAACCCACCTCTTCTTCGG | CGG | 55 | 26.99 | 136 | 12 | 1 | 149 | 69.487 | 3.471 | 1 |
| patec7-e1_f10 | CTCTTCTTCGGCGGCTATGC | AGG | 60 | 1.223 | 95 | 17 | 6 | 118 | 91.338 | 10.033 | 1 |
| patec7-e1_f11 | TCTTCTTCGGCGGCTATGCA | GGG | 55 | 49.707 | 29 | 20 | 1 | 50 | 98.028 | 47.886 | 1 |
| patec7-e1_f12 | TTCGGCGGCTATGCAGGGTA | AGG | 60 | 3.664 | 21 | 17 | 2 | 40 | 96.748 | 44.168 | 1 |
| patec7-e1_f13 | GGCTATGCAGGGTAAGGTGA | TGG | 55 | 19.99 | 53 | 58 | 3 | 114 | 89.751 | 13.093 | 1 |
| patec7-e1_f14 | CAAGCCGTCCCCAAGATCAA | TGG | 55 | 31.253 | 50 | 22 | 2 | 74 | 87.613 | 16.604 | 1 |
| patec7-e1_f15 | CCCCAAGATCAATGGCGCGA | AGG | 60 | 13.72 | 85 | 21 | 3 | 109 | 87.211 | 17.506 | 1 |
| patec7-e1_f16 | AAGATCAATGGCGCGAAGGT | TGG | 50 | 14.173 | 40 | 62 | 0 | 102 | 76.034 | 30.864 | 1 |
| patec7-e1_f17 | CCTGAAAGCTGAATCCCAAA | AGG | 45 | 63.511 | 156 | 90 | 0 | 246 | 77.872 | 9.329 | 1 |
| patec7-e1_f18 | AGCTGAATCCCAAAAGGCTG | AGG | 50 | 49.721 | 71 | 25 | 4 | 100 | 87.262 | 21.15 | 1 |
| patec7-e1_f19 | CTGCCCCTTCCTCAGCCCCG | AGG | 75 | 47.726 | 46 | 13 | 1 | 60 | 90.627 | 11.146 | 1 |
| patec7-e1_f20 | TCTATAATCAACTACCTGAC | TGG | 35 | 15.563 | 116 | 48 | 3 | 167 | 85.782 | 22.848 | 1 |
| patec7-e1_f21 | CGCCGTAACAACGATCTTTT | TGG | 45 | 5.074 | 11 | 16 | 1 | 28 | 97.941 | 73.918 | 1 |
| patec7-e1_f22 | TTTTGGCTGCCGAGAAGCAG | TGG | 55 | 58.582 | 230 | 108 | 2 | 340 | 63.833 | 4.322 | 1 |
| patec7-e1_f23 | AGCAGTGGACCCTTCTTGAT | TGG | 50 | 19.953 | 90 | 33 | 8 | 131 | 44.721 | 13.007 | 2 |
| patec7-e1_f24 | CCCTTCTTGATTGGAAGCCA | CGG | 50 | 39.574 | 117 | 73 | 0 | 190 | 58.155 | 9.343 | 1 |
| patec7-e1_f25 | CTTACTGATGCATTTAGCCT | TGG | 40 | 8.992 | 73 | 97 | 3 | 173 | 83.164 | 21.529 | 1 |
| patec7-e1_f26 | TTACTGATGCATTTAGCCTT | GGG | 35 | 35.915 | 190 | 498 | 2 | 690 | 69.198 | 2.815 | 1 |
| patec7-e1_f27 | TAGCCTTGGGAAGATTGTGC | AGG | 50 | 13.226 | 98 | 12 | 3 | 113 | 83.7 | 13.995 | 1 |
| patec7-e1_f28 | CTTGGGAAGATTGTGCAGGA | TGG | 50 | 22.511 | 119 | 42 | 1 | 162 | 81.952 | 10.908 | 1 |
| patec7-e1_f29 | TGCAGGATGGACTAGTTTTC | AGG | 45 | 10.455 | 88 | 47 | 4 | 139 | 45.47 | 16.536 | 1 |
| patec7-e1_f30 | TCAGGCAGAACTTTTCCATC | AGG | 45 | 36.873 | 140 | 51 | 3 | 194 | 66.554 | 10.058 | 1 |
| patec7-e1_f31 | TCCATCAGGTCATATGAGAT | TGG | 40 | 2.506 | 212 | 326 | 12 | 550 | 55.36 | 10.866 | 1 |
| patec7-e1_f32 | CCATCAGGTCATATGAGATT | GGG | 40 | 6.912 | 382 | 178 | 3 | 563 | 46.688 | 2.814 | 1 |
| patec7-e1_f33 | CATCAGGTCATATGAGATTG | GGG | 40 | 15.244 | 888 | 148 | 1 | 1037 | 16.502 | 1.914 | 1 |
| patec7-e1_f34 | CATATGAGATTGGGGCTGAT | CGG | 45 | 35.631 | 377 | 48 | 2 | 427 | 64.937 | 5.084 | 1 |
| patec7-e1_f35 | TGAGATTGGGGCTGATCGGA | CGG | 55 | 12.565 | 127 | 62 | 1 | 190 | 69.962 | 5.775 | 1 |
| patec7-e1_r1 | CCCAATCTCATATGACCTGA | TGG | 45 | 22.127 | 270 | 89 | 3 | 362 | 55.193 | 3.133 | 1 |
| patec7-e1_r2 | CATCCTGCACAATCTTCCCA | AGG | 50 | 21.62 | 85 | 32 | 2 | 119 | 81.475 | 15.989 | 1 |
| patec7-e1_r3 | AAATGCATCAGTAAGCATGT | CGG | 35 | 16.099 | 160 | 241 | 4 | 405 | 57.242 | 7.733 | 1 |
| patec7-e1_r4 | AATGCATCAGTAAGCATGTC | GGG | 40 | 9.93 | 113 | 307 | 1 | 421 | 54.877 | 16.368 | 1 |
| patec7-e1_r5 | GTAAGCATGTCGGGACGCCG | TGG | 65 | 85.368 | 7 | 47 | 0 | 54 | 90.921 | 45.019 | 1 |
| patec7-e1_r6 | GCCGTGGCTTCCAATCAAGA | AGG | 55 | 18.679 | 83 | 30 | 4 | 117 | 72.946 | 11.739 | 1 |
| patec7-e1_r7 | CCGTGGCTTCCAATCAAGAA | GGG | 50 | 56.44 | 116 | 41 | 1 | 158 | 60.138 | 14.417 | 1 |
| patec7-e1_r8 | AAGAAGGGTCCACTGCTTCT | CGG | 50 | 23.937 | 99 | 67 | 6 | 172 | 85.852 | 18.179 | 1 |
| patec7-e1_r9 | AGCCAAAAAGATCGTTGTTA | CGG | 35 | 5.249 | 210 | 68 | 1 | 279 | 77.013 | 9.706 | 1 |
| patec7-e1_r10 | CAAAAAGATCGTTGTTACGG | CGG | 40 | 15.313 | 68 | 45 | 4 | 117 | 59.687 | 8.915 | 2 |
| patec7-e1_r11 | AGATCGTTGTTACGGCGGCA | AGG | 55 | 19.236 | 9 | 13 | 7 | 29 | 97.941 | 50.135 | 1 |
| patec7-e1_r12 | GCAAGGAGCACGCTCCAGTC | AGG | 65 | 23.715 | 26 | 45 | 5 | 76 | 59.656 | 31.265 | 2 |
| patec7-e1_r13 | AGTTGATTATAGAATGTCCT | CGG | 30 | 26.389 | 197 | 178 | 3 | 378 | 73.428 | 9.066 | 1 |
| patec7-e1_r14 | GTTGATTATAGAATGTCCTC | GGG | 35 | 3.355 | 148 | 57 | 5 | 210 | 78.932 | 7.357 | 1 |
| patec7-e1_r15 | TTGATTATAGAATGTCCTCG | GGG | 35 | 66.595 | 182 | 57 | 2 | 241 | 80.596 | 17.045 | 1 |
| patec7-e1_r16 | ATAGAATGTCCTCGGGGCTG | AGG | 55 | 31.024 | 25 | 22 | 3 | 50 | 93.224 | 34.512 | 1 |
| patec7-e1_r17 | AATGTCCTCGGGGCTGAGGA | AGG | 60 | 13.141 | 63 | 16 | 6 | 85 | 87.968 | 16.124 | 1 |
| patec7-e1_r18 | ATGTCCTCGGGGCTGAGGAA | GGG | 60 | 25.615 | 45 | 16 | 6 | 67 | 88.619 | 20.768 | 1 |
| patec7-e1_r19 | TGTCCTCGGGGCTGAGGAAG | GGG | 65 | 1.923 | 396 | 26 | 1 | 423 | 64.75 | 5.706 | 1 |
| patec7-e1_r20 | CAGCATCTTCCTCAGCCTTT | TGG | 50 | 3.496 | 186 | 68 | 2 | 256 | 73.731 | 8.467 | 1 |
| patec7-e1_r21 | AGCATCTTCCTCAGCCTTTT | GGG | 45 | 3.44 | 385 | 76 | 0 | 461 | 55.289 | 2.5 | 1 |
| patec7-e1_r22 | CCTTTTGGGATTCAGCTTTC | AGG | 45 | 1.072 | 217 | 30 | 2 | 249 | 69.271 | 5.987 | 1 |
| patec7-e1_r23 | AACCTTCGCGCCATTGATCT | TGG | 50 | 0.769 | 32 | 24 | 2 | 58 | 56.739 | 37.152 | 1 |
| patec7-e1_r24 | ACCTTCGCGCCATTGATCTT | GGG | 50 | 16.572 | 35 | 44 | 4 | 83 | 58.455 | 22.164 | 1 |
| patec7-e1_r25 | CCTTCGCGCCATTGATCTTG | GGG | 55 | 8.279 | 82 | 35 | 3 | 120 | 87.294 | 16.103 | 1 |
| patec7-e1_r26 | CGCGCCATTGATCTTGGGGA | CGG | 60 | 27.97 | 31 | 18 | 2 | 51 | 86.164 | 17.599 | 1 |
| patec7-e1_r27 | CATTGATCTTGGGGACGGCT | TGG | 55 | 4.631 | 50 | 18 | 5 | 73 | 91.792 | 18.772 | 1 |
| patec7-e1_r28 | ATTGATCTTGGGGACGGCTT | GGG | 50 | 9.14 | 64 | 46 | 0 | 110 | 91.877 | 23.118 | 1 |
| patec7-e1_r29 | CTGCATAGCCGCCGAAGAAG | AGG | 60 | 22.46 | 79 | 29 | 5 | 113 | 83.833 | 15.01 | 1 |
| patec7-e1_r30 | CATAGCCGCCGAAGAAGAGG | TGG | 60 | 21.522 | 149 | 70 | 4 | 223 | 58.285 | 4.476 | 1 |
| patec7-e1_r31 | ATAGCCGCCGAAGAAGAGGT | GGG | 55 | 84.26 | 95 | 22 | 0 | 117 | 72.614 | 6.843 | 1 |
| patec7-e1_r32 | CGCCGAAGAAGAGGTGGGTT | TGG | 60 | 1.978 | 121 | 29 | 3 | 153 | 76.495 | 11.38 | 1 |
| patec7-e1_r33 | ACCCCGAACATCCAAATTCT | CGG | 45 | 4.115 | 74 | 73 | 3 | 150 | 83.578 | 20.793 | 1 |
| patec7-e1_r34 | ATTCTCGGAGCCTTCACCAA | TGG | 50 | 22.899 | 76 | 24 | 3 | 103 | 83.059 | 16.37 | 1 |
| patec7-e1_r35 | CGAAGCTTTTGCTGATGCCG | TGG | 55 | 11.71 | 86 | 15 | 5 | 106 | 45.953 | 10.659 | 1 |
| patec7-e1_r36 | GAAGCTTTTGCTGATGCCGT | GGG | 50 | 4.285 | 116 | 41 | 4 | 161 | 80.664 | 6.13 | 1 |
| patec7-e1_r37 | AAGCTTTTGCTGATGCCGTG | GGG | 50 | 66.525 | 380 | 57 | 3 | 440 | 46.168 | 5.031 | 1 |
| patec7-e1_r38 | AGCTTTTGCTGATGCCGTGG | GGG | 55 | 13.087 | 122 | 39 | 3 | 164 | 79.515 | 8.82 | 1 |
| patec7-e1_r39 | GATGCCGTGGGGGAGAAAGA | TGG | 60 | 43.192 | 332 | 103 | 1 | 436 | 49.512 | 3.396 | 1 |
| patec7-e1_r40 | CGTGGGGGAGAAAGATGGTG | TGG | 60 | 13.872 | 291 | 45 | 9 | 345 | 56.656 | 2.832 | 1 |
| patec7-e1_r41 | GTGGGGGAGAAAGATGGTGT | GGG | 55 | 8.669 | 378 | 48 | 7 | 433 | 51.328 | 3.1 | 1 |
| patec7-e1_r42 | TGGGGGAGAAAGATGGTGTG | GGG | 55 | 13.342 | 340 | 78 | 13 | 431 | 59.066 | 5.199 | 1 |
| patec7-e1_r43 | GAAAGATGGTGTGGGGAAAA | AGG | 45 | 0.918 | 351 | 62 | 8 | 421 | 58.351 | 2.608 | 1 |
| patec7-e1_r44 | TGTGGGGAAAAAGGCCGAAG | CGG | 55 | 34.519 | 130 | 23 | 1 | 154 | 72.319 | 10.713 | 1 |
